# Pleistocene climate change and the formation of regional species pools

**DOI:** 10.1101/149617

**Authors:** Joaquín Calatayud, Miguel Ángel Rodríguez, Rafael Molina-Vengas, María Leo, Jose Luís Hórreo, Joaquín Hortal

## Abstract

This preprint has been reviewed and recommended by Peer Community In Evolutionary Biology (https://doi.org/10.24072/pci.evolbiol.100053). Despite the description of bioregions dates back from the origin of biogeography, the processes originating their associated species pools have been seldom studied. Ancient historical events are thought to play a fundamental role in configuring bioregions, but the effects of more recent events on these regional biotas are largely unknown. We use a network approach to identify regional and sub-regional faunas of European *Carabus* beetles, and analyse the effects of dispersal barriers, niche similarities and phylogenetic history on their configuration. We identify a transition zone matching the limit of the ice sheets at Last Glacial Maximum. While southern species pools are mostly separated by dispersal barriers, in the north species are mainly sorted by their environmental niches. Strikingly, most phylogenetic structuration of *Carabus* faunas occurred during the Pleistocene. Our results show how extreme recent historical events –such as Pleistocene climate cooling, rather than just deep-time evolutionary processes, can profoundly modify the composition and structure of geographic species pools.

## Introduction

Naturalists have long been captivated by the geographic distribution of world biotas. Rooted in the seminal ideas by Alexandre von Humbolt, this fascination has promoted a long-term research agenda aiming to delineate biogeographic regions according to their integrating faunas and floras (e.g. Wallace 1876, Holt et al. 2013, Rueda et al. 2013). Besides this, the large-scale eco-evolutionary processes that shape regional biotas are known to influence ecological and evolutionary dynamics at finer scales (Ricklefs 2008, 2015). For instance, regional species pools can modulate local diversity patterns (Ricklefs 2011, Medina et al. 2014, Ricklefs and He 2016), the structure and functioning of ecosystems (Naeslund and Norberg 2006), or co-evolutionary processes (Calatayud et al. 2016a). However, despite their fundamental importance, the processes that have configured regional biotas have been seldom studied (and particularly the historical ones), and most explanations on their origin and dynamics remain largely narrative (Crisp et al. 2011).

Perhaps the earliest speculations about the formation of regional species pools took place during the flourishment of bioregionalizations in the mid-19th century (reviewed by Ebach 2015). During that time, and beyond referring to geophysical factors (climate, soils, and physical barriers), some authors already started to emphasize historical influences as key elements determining the configuration of plant and animal regions. For instance, when Wallace (1876) proposed his ground-breaking zoogeographic regions, he argued that while the distribution of ancient linages such as genera and families would likely reflect major geological and climatic changes spanning the early and mid-Cenozoic, species distributions would be more influenced by recent events such as Pleistocene glaciations (see Rueda et al. 2013). These recent events could have fostered many additions and subtractions of species to regional faunas through dispersal and diversification processes. Indeed, increasing evidence suggests that Pleistocene glacial-interglacial dynamics may have driven population extinctions (e.g. Barnes et al. 2002), allopatric speciation in glacial refugia (e.g. Johnson et al. 2004) and post-glacial recolonization events (e.g. Hewitt 1999; Theissinger et al. 2013). Besides shaping phylogeographic patterns (Avise et al. 1998; Ursenbacher et al. 2006; Sommer and Nadachowski 2006; Provan and Bennett 2008), all these processes are likely underpinning diversity patterns for many taxa, particularly in the Holarctic (e.g. Svenning and Skov 2007; Hortal et al. 2011; Calatayud et al. 2016b). However, whether the signature of Pleistocene glaciations scales up to the configuration of regional biotas remains largely unknown.

Historical contingencies should act over the intricate interplay between ecological (i.e. environmental tolerances and dispersal) and evolutionary (i.e. diversification and adaptation to new habitats) processes underpinning the composition and structure of regional species pools. On the one hand, niche-based processes may determine the composition of regional species pools (Mittelbach and Schemske 2015), mainly throughout their effects on species distribution ranges (Soberon 2007, Hortal et al. 2010, 2012). These processes integrate responses to abiotic conditions along geographical gradients and to local and regional biotic environments (Colwell et al. 2009), which may ultimately lead to the appearance of distinct regional communities in areas of contrasted environmental conditions (Ricklefs 2015). Although species with similar environmental tolerances/preferences can coexist in regions of similar climate, their dispersal may be constrained by geographical barriers, which may lead to divergent species pools under similar environmental conditions. Finally, evolutionary processes also constrain all these mechanisms. For instance, environmental-driven regions may be expected if occupancy of new areas is constrained by niche conservatism (Hortal et al. 2011), which should also lead to species pools integrated of evolutionary related species (i.e. niche conservatism generating phylogenetically clustered species pools, Fig.1a). This, however, can be in turn filtered by biogeographical processes (Gouveia et al. 2014). As such, diversification of lineages within regions separated by strong dispersal barriers (e.g. mountain ranges) may also lead to phylogenetically clustered pools of locally adapted species (i.e. geographically driven niche conservatism; Fig.1a; Warren et al. 2014, Calatayud et al. 2016a). Historical contingencies may contribute to the configuration of regional pools by modifying the balance between these processes. For example, differential diversification rates may be the predominant driver of regional species pools during climatically stable periods (Cardillo 2011). Yet, regions with a greater influence of climatic fluctuations such as Pleistocene glaciations may harbour pools of species mostly shaped by the joint effects of current climate and post-glacial colonization dynamics (Svenning et al. 2015), as well as by species’ competition during these colonization processes (Horreo et al. 2018), thus eroding the signature of geographically-structured diversification processes.

In this study we aim to disentangle the relative importance of the processes that may contribute to the formation of regional species pools, using European *Carabus* (Coleoptera: Carabidae) as a model lineage. *Carabus* is a species-rich ground beetle genus of great popularity due to the beautiful jewel-like appearance of some species (Turin et al. 2003). In general, *Carabus* species are flightless nocturnal predators of snails, earthworms and caterpillars. They hold hydrophilic adaptations and are typically associated to deciduous forests (Deuve et al. 2012). Previous evidence suggests that the richness of species from this genus in Europe is determined to a large extent by both current environmental conditions (i.e. climate and habitat) and glacial-interglacial dynamics (Calatayud et al. 2016b). This makes European *Carabus* an ideal case study to evaluate the joint effects of evolutionary, ecological and historical contingency processes as drivers of regional species pools.

**Figure 1.**
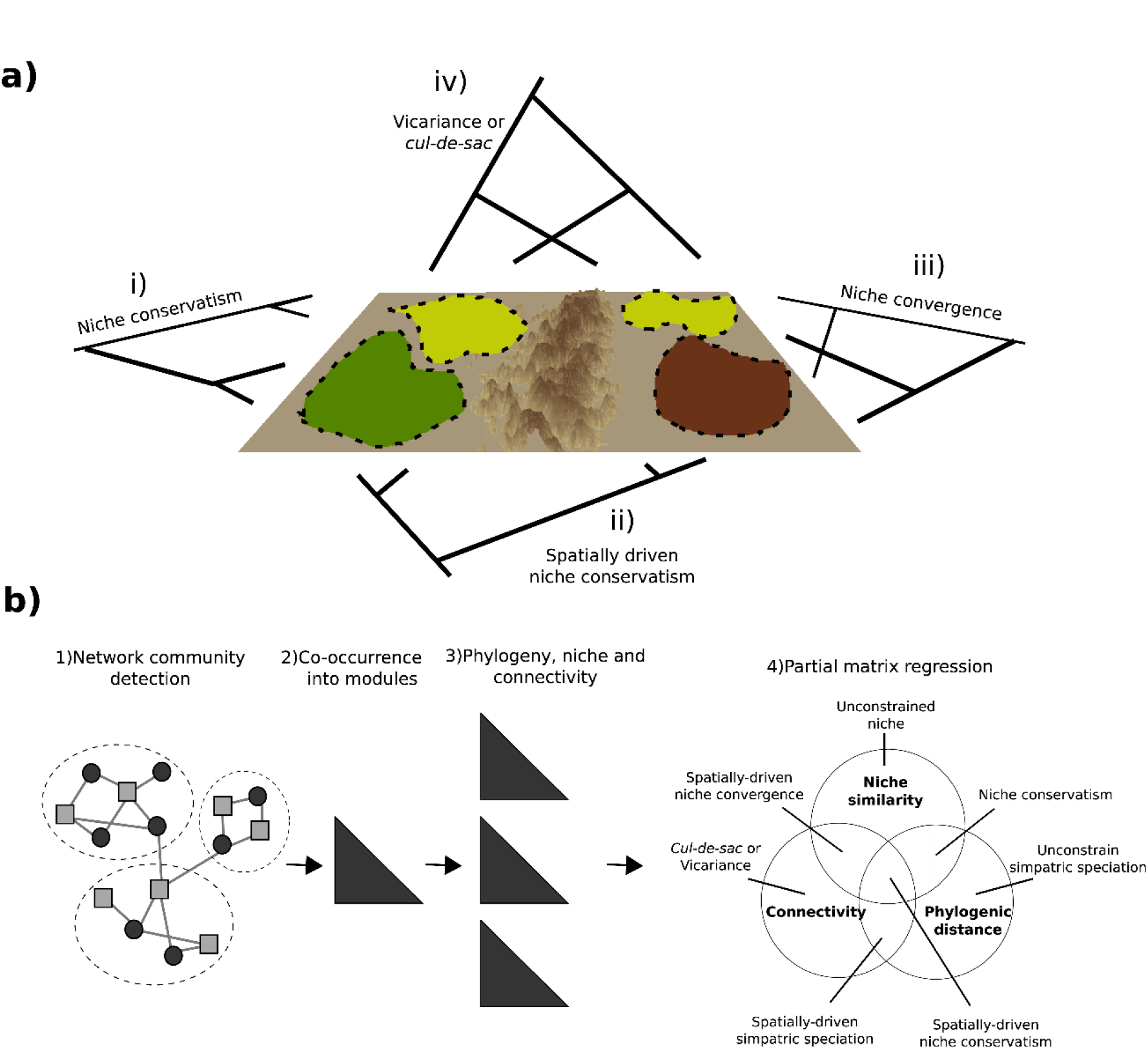
Identifying the factors configuring regional faunas. a) Four (out of seven, see b) hypothetical processes that may configure Regional faunas. Dotted lines depict different regions while colours correspond with different climates. In each case, the tips of the phylogeny point to regional distribution of the species. b) Workflow and potential results: 1) Hypothetical results of modularity analysis over the occurrence network; 2) similarity matrix of occurrence into modules; 3) pairwise matrix of environmental niche similarities; phylogenetic distances and topographical connectivity; and 4) hypothetical results and interpretations of a partial matrix regression on species occurrence similarities as a function of niche similarities, phylogenetic distances and connectivity.

Specifically, we use data on the distribution and evolutionary relationships of *Carabus* species, along with network and phylogenetic analyses, to evaluate six hypotheses: First, given the presumed low dispersal capacity of the species from this genus (Turin et al. 2003), we hypothesize that (H1) European *Carabus* species pools are mainly shaped by the main orographic barriers of the continent, but also, that (H2) glacial-interglacial dynamics have led to strong differentiation between northern and southern regional species pools. If this differentiation is true, northern European *Carabus* faunas will be comprised of species that colonized newly vacant habitats after the withdrawal of the ice sheet, and hence (H3) their regional distribution will be mostly determined by current climate. In contrast, (H4) southern faunas will be mainly shaped by the joint influence of diversification events and dispersal limitations, due to the combined effect of higher climatic stability (e.g. climatic refugia) and a more complex orography (Alps, Pyrenees, Carpathians). Therefore, (H5) species forming northern regional pools will exhibit comparatively lower levels of regional endemicity, whereas those forming southern regional pools will show comparatively higher levels of regional affinity. Finally, according to Wallace (1876), the advance and retreat of the ice sheets during the Pleistocene should have determined the spatial distribution of lineages, eroding the effects of the former configuration of the distribution of the main *Carabus* lineages. Therefore, (H6) we expect a temporal signal coincident with the Pleistocene in the phylogenetic structure of *Carabus* faunas, and no effect of deep-time events on the current geographical distribution of these lineages.

## Material and methods

### Rationale and structure of the analyses

Exploring the determinants of regional faunas requires jointly analysing ecological, evolutionary and historical factors. We did so through three consecutive steps (Fig.1b). First, we identified distinct regional species pools within Europe by using a network community detection algorithm. From this analysis we derived a species pairwise similarity matrix of occurrence into different modules, each one representing different regions. Second, we assessed the relative importance of environmental, spatial and evolutionary determinants of such similarity. To do so, we constructed four pairwise matrices to describe ecological, topographical and evolutionary relationships among species; namely, *i*) a matrix of climatic niche similarity, *ii*) a matrix of habitat similarity, *iii*) a matrix of spatial connectivity among distributional ranges, and *iv*) a phylogenetic distance matrix. Then, we used generalized partial matrix regressions to model the similarity in species occurrences as a function of these four matrices (Fig.1b). We used this workflow to explore the factors involved in the configuration of *Carabus* faunas both at regional (i.e. through analysing *Carabus* species co-occurrence across regions) and sub-regional scale (i.e. focusing on co-occurrence patterns within sub-regions). Finally, we also applied ancestral range estimation analysis to identify the time period from which ancestral areas are estimated with less uncertainty. By doing so, we aimed to detect important historical periods contributing to the regional organization of *Carabus* lineages.

The interpretation of the joint and independent effects of explanatory matrices can shed light on the different processes configuring regional faunas (see Fig.1a). Thus, if niche similarities (i.e. represented by the climatic and habitat similarity matrices) and phylogenetic distances altogether explained the regional co-occurrence of species, then this could be interpreted as indicative of constrained niche evolution (or a tendency to resemble ancestral niches) in shaping regional faunas (Fig.1a.i). However, if spatial connectivity also accounted for part of this co-occurrence, this would indicate that this niche conservatism pattern can be caused by geographical constrains (Fig.1a.ii). Further, the effects of niche similarities and spatial connectivity alone (i.e. without phylogenetic signal) can be most likely the consequence of a convergence of climatic niches due to geographic isolation (Fig.1a.iii), whereas the effects of connectivity and phylogeny would be indicative of a primacy of intra-regional speciation driven by geographical barriers. Niche similarities alone would point to an unconstrained niche evolution shaping regional faunas, while phylogeny alone would indicate a primacy of geographically unconstrained intra-regional speciation events. Finally, either a *cul-de-sac* effect (i.e. the accumulation of species in past climatic refugia) or a primacy of vicariant speciation events could lead to the existence of independent effects of connectivity and regional co-occurrence (Fig.1a.iv).

### Identification of regional species pools

We took advantage of community detection analysis —borrowed from network theory— to identify *Carabus* regional species pools in Europe. We first generated a bipartite network where species and grid cells constitute two disjoint sets of nodes that are connected according to the presence of species in grid cells (e.g. Calatayud et al. 2016a). The species presence data comes from expert-based range maps of all *Carabus* species inhabiting Europe (n = 131, Turin et al. 2003) overlaid into a 100-km equal-area grid based on the LAEA pan-European grid system (available at https://inspire.ec.europa.eu/, see Calatayud et al. 2016b for details). Then, we conducted a modularity analysis using the index proposed by Barber (2007) and the Louvain algorithm (Blondel et al. 2008) as implemented in the Matlab function “Gen Louvain”, (available at http://netwiki.amath.unc.edu; Mucha et al. 2010). This analysis is intended to find groups of nodes (i.e. species and grid cells) that are more densely connected. Hence, in our case, the analysis identified groups of grid cells, each group sharing *Carabus* species mainly distributed within its cells (i.e. regions and their associated faunas). The Louvain algorithm was run 100 times, and the network partition showing highest modularity value was retained. This optimal solution was used to conduct all subsequent analyses, although all the solutions were qualitatively similar. We evaluated the statistical significance of the modules by comparing their associated modularity value to a null distribution of values (n = 100) where the original presence-absence matrix was randomized using the independent swap algorithm (a fixed-fixed null model implemented in the R package “picante”, Kembel et al. 2010). Finally, to detect potential sub-modules (i.e. sub-regions) nested within modules (i.e. sub-regional species pools within regional species pools), we derived a new bipartite network from each of the previously identified modules, and applied the procedure described above in each case.

It is important to note that despite species and grid cells were assigned to just one module, they could also occur in other modules with different degrees of specificity. For example, despite most species in a grid cell will belong to the same module the cell does, this cell could also hold species that are primarily associated to other modules. Similarly, although a species will mostly be present in cells assigned to its module, it may also occur in cells from other modules. Thus, we calculated the degree of module specificity for each node (i.e. species and grid cells) as its number of links with nodes of its module divided by its total number links (see Guimera and Amaral’s 2005 inter-modular participation index for a similar metric). Thus, higher module specificity would correspond to species mainly distributed within its module (highly endemic species), as well as to cells pertaining to well-defined regions; whereas lower module specificity would indicate widespread species and cells located in transition zones.

### Assessing the determinants of regional species pools

To disentangle the determinants of the current configuration of *Carabus* faunas in Europe, we first generated a species-per-module matrix, where each entry of the matrix represents the percentage of the distributional range of a certain species that lies in a given module. Then, we derived a co-occurrence pairwise similarity matrix from the former matrix using the Schoener’s index (Schoener 1970). This metric quantifies the overlap between species pairs throughout the modules (see Krasnov et al. 2012 for a previous application) and it ranges from 0 (no overlap) to 1 (identical distribution across modules). Note that this similarity matrix reflects the co-occurrence similarities at regional scale, thus ignoring distributional patterns at lower spatial scales (i.e. two species may have identical regional distribution but appear as segregated at the local scale). The resultant co-occurrence pairwise similarity matrix was used as dependent variable. We generated four different pairwise dis/similarity matrices to be used as explanatory variables. Two of them were used to account for environmental factors: (i) a climatic-niche similarity matrix and (ii) a habitat similarity matrix. The remaining two considered geographical and evolutionary factors: (iii) a spatial-connectivity matrix and (iv) a phylogenetic distance matrix.

i. *Climatic-niche similarity matrix*. We characterized the climatic niche of each *Carabus* species in the dataset following a similar approach as proposed by Broennimann et al. (2012). We selected six bioclimatic variables to account for the main water and energy aspects of climate – namely mean annual temperature, temperature of the warmest quarter, temperature of the driest quarter, total annual precipitation, total precipitation of the warmest quarter and total precipitation of the driest quarter– and altitudinal range to account for the effects of mesoclimatic gradients within each grid cell. These variables may be among the main determinants of the distribution of *Carabus* species diversity within Europe (see Calatayud et al. 2016b). Bioclimatic variables were extracted from Worldclim (v1.4 Hijmans et al. 2005; available at http://www.worldclim.org/), whereas altitudinal data were derived from the 30-arcsecond digital elevation model GTOPO30 (available at https://lta.cr.usgs.gov/GTOPO30). We conducted a principal component analysis on these variables to obtain a bidimensional climatic space defined by the two main axes, that explained 81.4% of the variance (Fig. S1). Finally, we divided this climatic space into a 100×100 grid and calculated species overlap in the gridded space using Schoener’s index (see above).
ii. *Habitat similarity matrix*. The distribution of *Carabus* species may also be shaped by forest preferences (Turin et al. 2003). Accordingly, we used ten vegetation categories derived from MODIS Land Cover at 5-minute resolution (Evergreen broadleaf forest, deciduous needle-leaf forest, deciduous broadleaf forest, mixed forest, closed shrub lands, open shrub lands, woody savannas, savannas and grasslands; Channan et al. 2014, available at http://glcf.umd.edu/data/lc/). For each species we computed the proportion of each category overlaying its range. With this, we computed pairwise similarities in the preference for different vegetation types using Schoener’s index (see above).
iii. *Spatial connectivity matrix*. To evaluate the potential influence of geophysical barriers to dispersal on the current distribution of *Carabus* species, we first created a dispersal-cost surface by dividing the study area in 1 Km^2^ grid cells and weighting each cell according to its topography (in this case, slope) and the presence of water bodies. Slope values ranged from 0 to 100 at each pixel, being 0 the lowest dispersal cost and 100 the highest one, and were determined from GTOPO30 altitudinal data using the GRASS tool r.slope (GRASS Development Team 2017). Grid cells including water bodies as rasterized layers in the Nature Earth database (available at http://www.naturalearthdata.com/) were further weighted by assigning arbitrary values of friction to the dispersal of *Carabus* species, namely 30% for cells containing rivers and lakes and 99% for cells that lay on sea water masses (note that *Carabus* species show hydrophilic adaptations). Then, the connectivity between all pairs of cells was calculated as the least-cost path over the dispersal-cost surface that connects both cells, using the “gdistance” R package (van Etten 2015). Finally, the spatial connectivity between each pair of species’ distributional ranges in the dataset was estimated as the average distance among all grid cells within the range of each species. Average distances were preferred over absolute least-cost distances to avoid disproportionate differences in spatial connectivity between overlapping and non-overlapping distributional ranges.
iv. *Phylogenetic distance matrix*. To unravel the evolutionary history of the *Carabus* lineage and assess the potential importance of evolutionary processes in determining the formation of *Carabus* species pool, we reconstructed a species-level time-calibrated molecular phylogeny including the 89 species for which we found available DNA information on ten markers (eight mitochondrial regions: 12S rDNA,16S rDNA, 28S rDNA, ND4, ND5, COI, Cytb and PEPCK; plus two nuclear ones: anonymous locus and wingless; see Tables S1 and S2). We aligned each marker independently using different algorithms: MAFFT (Katoh and Standley 2013), Clustal X (Thompson et al. 1994; Larkin et al. 2007), MUSCLE (Edgar 2004) and Kalign (Lassmann and Sonnhammer 2005), and selected the most reliable alignment for each marker using the multiple overlap score (MOS) provided by MUMSA (Lassmann and Sonnhammer 2006). We removed ambiguous or poorly aligned positions from the alignments with trimAl (Capella-Gutiérrez et al. 2009). The final dataset was concatenated in 5603 basepairs following a total-evidence approach (Kluge 1998), and was used to conduct Bayesian phylogenetic inference with BEAST v.2.4.6 software (Bouckaert et al. 2014). We used a GTR model for sequence evolution, a Random Local Clock, a birth-death model prior, and 100 million of MCMC chain length searching for convergence. There is ongoing debate on the divergence time of the genus *Carabus* (Andujar et al. 2012, Deuve et al. 2012), hence molecular dating was conducted under two different scenarios. Firstly, according to Deuve et al.’s (2012) molecular dating, the crown age of *Carabus* was set at 17.3 Mya. Secondly, and according to Andujar et al. (2012), the origin of the group was set at 25.16 Mya. We used the software Tracer v 1.6 (available at http://tree.bio.ed.ac.uk/software/tracer/) to check for MCMC chains convergence, and FigTree v.1.4.2 (available at http://tree.bio.ed.ac.uk/software/figtree/) for viewing and editing the phylogenetic trees (see Figures S2 and S3). To account for topological and time-calibration uncertainties, we used 100 phylogenies sampled from the posterior distribution for each molecular dating scenario. In addition, we used taxonomic information and phylogenetic uncertainty methods (Rangel et al. 2015) to place species lacking molecular information into the phylogeny (see Appendix S1). Thus, we derived 100 different phylogenetic hypotheses from each Bayesian posterior phylogeny by randomly inserting missing species within their most derived consensus clade based on taxonomic knowledge. In total, we generated 20,000 phylogenetic hypotheses (100 phylogenies per two molecular dating scenarios per 100 phylogenies accounting for uncertainties associated with lack of molecular data) that were used in subsequent analyses. Pairwise phylogenetic distances were calculated for each calibrated phylogeny using the function *cophenetic* implemented in the APE R package (Paradis et al. 2004).

We used generalized multiple regression on distance matrices and deviance partitioning to disentangle the relative importance of climatic niche, habitat preferences, dispersal barriers and evolutionary history in determining *Carabus* species pools in Europe. First, we conducted single regressions between the co-occurrence pairwise similarity matrix and each of the four explanatory matrices described above to seek for significant associations between the variables. We set a binomial family for error distribution and “logit” as the link function (see Ferrier et al. 2007 and Calatayud et al. 2016a for a similar approach). To assess for significance, we randomized the observed species per module matrix using the independent swap algorithm (see above) to derive 999 null occurrence similarity matrices. Then, we used simple regressions to relate each null similarity matrix with each one of the explanatory matrices. The relationship between an explanatory matrix and the observed species per module matrix was considered to be significant when it explained a higher proportion of the deviance than 99% of the regressions performed on the null matrices. In the case of phylogenetic pairwise distances we repeated this procedure for each phylogenetic hypothesis to consider phylogenetic uncertainties, applying the same criterion for significance. Finally, we retained those variables that showed significant relationships, and conducted variance partitioning among explanatory matrices (Legendre and Legendre 2012) to explore patterns of covariation among niche similarities (i.e. climatic and habitat similarity matrices), dispersal barriers and phylogenetic history. We conducted the analyses for the co-occurrence into modules (i.e. regions) and sub-modules (i.e. sub-regions) both at a European and regional (i.e. co-occurrence into sub-modules of each module) scales.

### Ancestral range estimation

To assess whether deep historical signals were eroded by Pleistocene glaciations we used probabilistic models of geographic range evolution. These models estimate ancestral range state probabilities assuming different processes involved in range evolution. We used the Dispersal–Extinction–Cladogenesis model of range evolution (DEC; Ree and Smith 2008) implemented in the R package BioGeoBears (Matzke 2014) since this model has been shown to perform well even under complex scenarios (Beeravolu and Condamine 2018). Species ranges were coded as present/absent in each module detected in the former network clustering. We used this analysis for 1,000 randomly selected phylogenies of each dataset, since preliminary results showed little variations among phylogenies. The estimation of ancestral ranges usually tends to be more ambiguous in deeper nodes of the phylogeny, as the lability of geographical ranges would tend to blur deep-time signals (Losos and Glor 2003). Similarly, if the Pleistocene glacial periods had important effects on species distributions it could be expected that ancestral range estimations will increase in accuracy around the Pleistocene. That is, pre-Pleistocene signals on the evolution of species distribution ranges will be blurred. To explore this, we evaluated the existence of changes in the relationship between node age and the marginal probability of the single most-probable ancestral state at each internal node. Then, general additive mixed models (GAMMs) were fitted to the node marginal probability as a function of node age, including the phylogenetic hypothesis as a random factor. We also used generalized linear mixed models (GLMMs) combined with piecewise regression to detect potential major breakpoints (i.e. temporal shifts) in the relationship between marginal probability and node age for each of the two dating scenarios. Given the large amount of observations (130 nodes x 1,000 phylogenies) we firstly estimated the breakpoint independently for each phylogenetic hypothesis. To do so, we included the breakpoint as a new parameter in a generalized linear model of marginal probabilities as a function of node age, minimizing the deviance of the fitted model using the function “optimize” in the R package lme4 (Bates et al. 2014). Finally, to assess significance we used the averaged breakpoint value in a GLMM with the same fixed formulation but including the phylogenetic hypothesis as a random factor. Because node marginal probabilities ranged between 0 and 1, we used a binomial family and a loglink function to fit all models.

General assumptions of probabilistic ancestral range estimation models may compromise subsequent interpretations (Ree and Sanmartin 2018). Moreover, our dataset does not fulfil the assumption of ancestral range estimation models that phylogenies should include all extant lineages, since some Carabus linages have representatives from outside Europe (mainly Asiatic species) that were not included in the analyses. Hence, we conducted additional analyses to provide further evidence on the temporal signal in the phylogenetic structuration of *Carabus* faunas coinciding with Pleistocene glaciations. To do so, we first generated a binomial variable based on the module affiliation of the species descending from each internal phylogenetic node, coding the phylogenetic nodes whose all descendant species were classified into the same region as 1, and 0 otherwise. Then, we conducted the analytical approach explained above, in this case fitting this binomial variable as a function of node age. This analysis would further elucidate if there is a breakpoint in the relationship between node age and the probability that all descendant species belong to the same region (see Appendix S2 for details and another complementary approach). Finally, we also explored whether there are differences between regions in the Pleistocene signal on the phylogenetic structure of their faunas. For each region we calculated the probability that a phylogenetic node has all its descendant species within the region, independently for nodes occurring either before and after the beginning of the Pleistocene (2.59 Mya; herein pre-Pleistocene and post-Pleistocene nodes). This probability was calculated as the number of pre- or post-Pleistocene nodes having all descendants associated to region *i* divided by the total number of nodes in the phylogeny.

All analyses were carried out in R (R core team 2015), using the function *bam* of the package mcvg for GAMM (Wood at al. 2015) and the package Lme4 for GLMM analyses (Bates et al. 2014).

## Results

### Identification of regional faunas

The *Carabus* occurrence network was significantly modular (M=0.385, p=0.01), dividing Europe in seven modules that group zoogeographically distinct regions with their associated faunas (i.e., different regional species pools; Figs. 2a and S4). Furthermore, all modules but module 2 showed significant sub-modular structure, presenting a decrease in modularity with latitude (mean M=0.316, ranging from 0.154 to 0.468; all p-values < 0.05, see Table S4). Module 1 holds 21 species mainly living in South-western Palearctic (Iberian Peninsula, North of Africa, Balearic Islands, Corsica, Sardinia and the western half of Sicilia). This module was subdivided into four submodules. Module 2 included only two species, both endemic of Crete. Module 3 identified an East Mediterranean region including the Italic Peninsula, part of Greece and Turkey. This module holds 18 species and was subdivided into five submodules. Module 4 depicted a Central European region embracing the Alps and the Carpathian Mountains, as well as Central European plains. This module showed the highest species richness, including 49 *Carabus* species, and was split into four submodules. Module 5 and module 6 comprised northern regions and showed the lowest species richness values, holding 10 species each. The former comprised Iceland and the British Isles and extended eastward up to the vicinity of the Ural Mountains. The latter included this mountain range and expanded to the easternmost zone of the study area. Both modules were divided into 3 submodules. Finally, module 7 included 21 species and embraced a south-eastern central European region expanding from the Carpathian Mountains to the south Ural Mountains. This module was split into three submodules.

**Figure 2.**
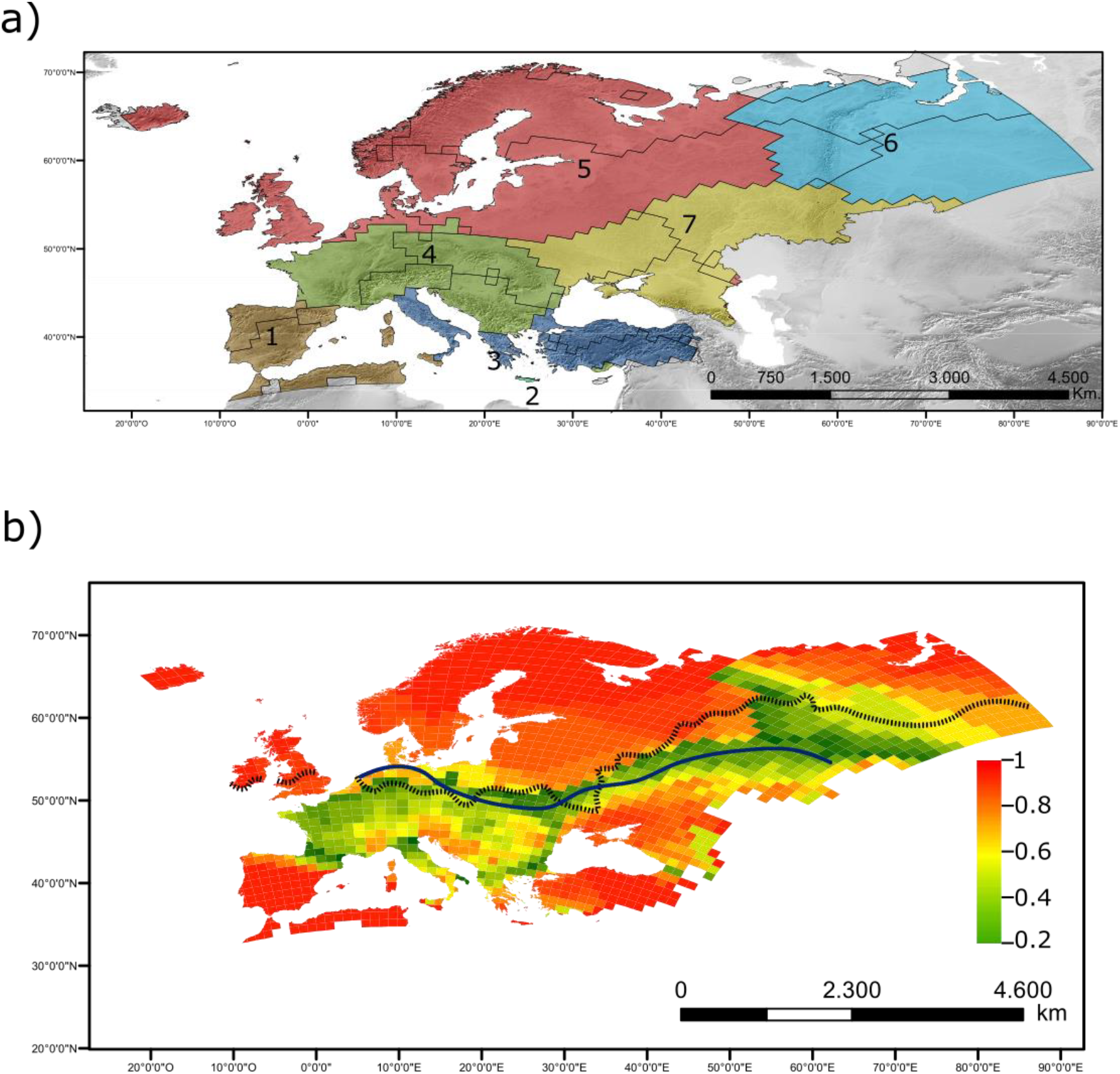
Transition zones between regions were associated to geophysical accidents and the border of the ice sheet at LGM. European *Carabus* regions found by the network community detection analysis. a) Geographical location of modules (i.e. regions) and submodules (i.e. sub-regions). b) Values of module affinity per grid cell; green colours (i.e. cells with low affinity) identify transition zones. The dotted black line corresponds with the southern limit of the ice sheet at LGM (extracted from Ehlers and Gibbard 2004). The blue line depicts the breakpoint where the temperature-Carabus richness relationship changes, as found in Calatayud et al. (2016b).

Regarding transition zones between regions, and in agreement with our first hypothesis, we found that they were clearly associated with geographical barriers such as the Pyrenees, the Alps, the Carpathian and the Ural Mountains, as well as the Turkish Straits System (that connects the Black Sea to the Aegean separating the Anatolian and Greek peninsulas; Fig. 2b). Interestingly, we also identified a west-to-east transitional belt between southern and northern regions that closely followed the southern limits of the ice sheet at the Last Glacial Maximum (LGM). This transitional zone further suggested a link between the configuration of *Carabus* regional faunas and Pleistocene glacial conditions, supporting our hypothesis 2.

### Correlates of regional co-occurrence

Matrix regressions showed that deviance of species co-occurrences across regions, across sub-regions and within each region was significantly explained (p<0.01), primarily by environmental niche similarity, and secondarily by spatial connectivity, except for northern regions (i.e. modules 5 and 6; Fig. 3 and Table S5). In contrast, relationships with evolutionary relatedness were non significant in all instances and regardless of the phylogenetic hypothesis used (p>0.01 in all cases, see Table S5). Comparing both types of subdivisions, environmental niche similarity explained more deviance across sub-regions than across regions, whereas spatial connectivity did the reverse (see Fig. 3). Comparing explained deviances between regions, the primacy of environmental niche similarity (mostly climate, see Table S5) in the northern ones (5 and 6) is consistent with the notion that northern regional pools are geographically sorted by current climate (our hypothesis 3), whereas the importance shown by spatial connectivity in the remaining regions is consistent with the more complex orography of central and southern Europe (consistent with hypothesis 4).

**Figure 3.**
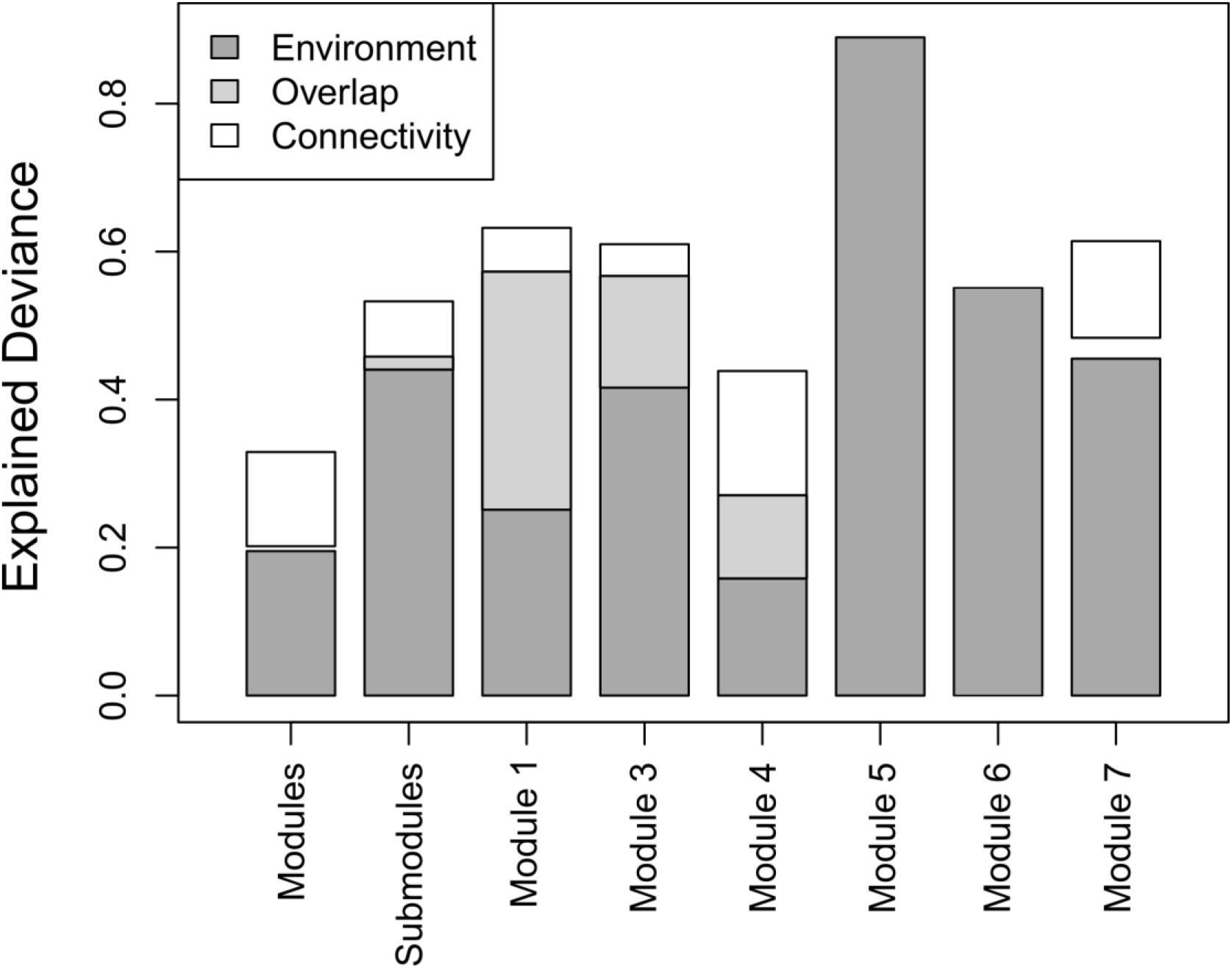
Regional co-occurrence was only explained by environmental niche similarities and topographical connectivity. Results of the partial generalized matrix regression of similarity in regional co-occurrence, as a function of environmental niche similarity (climate and habitat), topographical connectivity and phylogenetic distances. The first and second bars correspond with the models including co-occurrence similarities among all modules and submodules, respectively. The remaining bars correspond with the models where the similarities in submodule occurrence were analysed independently for the species of each module.

### Ancestral range estimation

Both phylogenetic datasets (i.e. alternative calibration scenarios) yielded similar qualitative and quantitative results (see Appendix S2). Thus, we only present here ancestral range estimations based on Deuve’s et al. (2012) calibration. GAMM results showed that node marginal probability of the most probable state increased towards younger nodes (P<0.01, explained deviance =7.49%, Fig. 4a). However, this increase showed a steep increment coinciding with the Pleistocene. Indeed, piecewise regression revealed that the relationship between marginal state probability and node age changed at 1.51 Mya (median value; with 45^th^ and 55^th^ percentiles at 1.24 and 1.89 Mya., respectively; Fig. 4a and S5), suggesting that most of the phylogenetic structuration of *Carabus* faunas began around the Pleistocene. Similarly, the probability for a phylogenetic node to have all descendant associated to same region increased toward younger nodes with a break point roughly associated with the Plio-Pleistocene transition (3.70 Mya.; 45^th^ and 55^th^ percentiles at 2.47 and 4.48 Mya.; see Fig. S6). In both cases the breakpoints associated to the Pleistocene were significant (P < 0.01).

**Figure 4.**
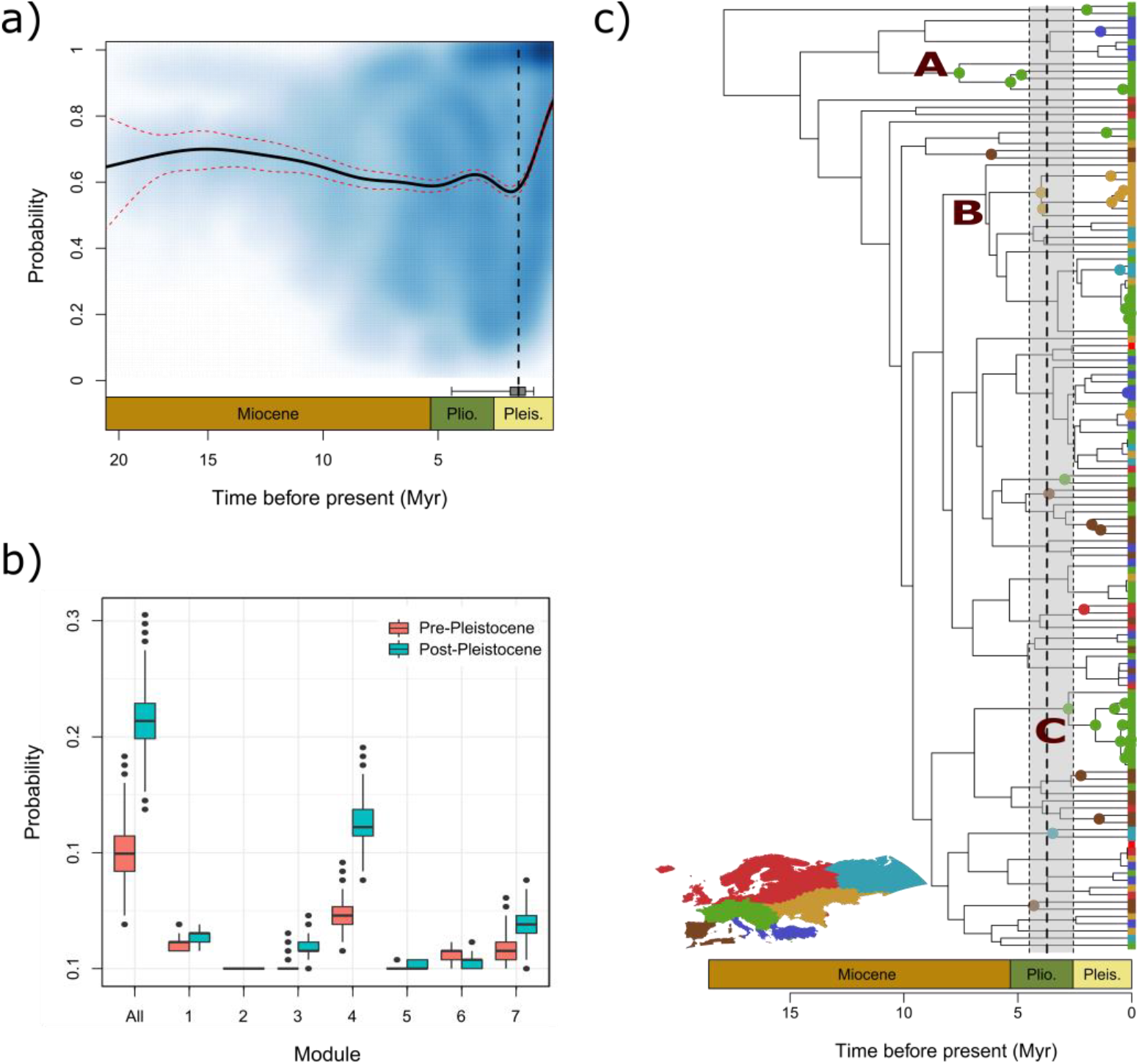
Temporal coincidence between the Pleistocene and the phylogenetic structuration of *Carabus* regions. a) GAMM predictions of the marginal probability of the most probable state as a function of node age. The dashed red lines correspond with the interval confidence at 95%. The dotted black line represents the median of the breakpoint found by piecewise GLM regressions. The boxplot at the bottom represent the 45^th^ and 55^th^ percentile breakpoint values, whereas the whiskers depict the 25^th^ and 75^th^ percentiles. b) Boxplot showing the probability of finding a phylogenetic node having all descendant species grouped in the same region for pre-Pleistocene (pink) and post-Pleistocene nodes (blue). This probability was calculated for all regions jointly (“All” in x axis) and independently (labelled according to Fig. 2a in x axis). c) An example of a phylogenetic hypothesis where internal nodes are coloured if all their descendant species grouped in the same region. Node and tip colours correspond to the regions where species were grouped, following Fig. 2a and the map at the bottom. Lineages showing a high number of related species belonging to the same region are highlighted as: “A” for *Platycarabus* subgenus; “B” for *Morphocarabus*; and “C” for *Orinocarabus*. The dashed line corresponds with the average breakpoint (median) where the probability of finding a node with all descendant species grouped in the same region increases (see also Fig. S6). The shaded area depicts the 45^th^ and 55^th^ percentiles.

In agreement with these results, we found that the probability of finding phylogenetic nodes having all descendant belonging to the same region was higher for post-Pleistocene nodes (median at 0.21; 25^th^ and 75^th^ percentiles at 0.20 and 0.23, respectively; Fig. 4a) than for pre-Pleistocene ones (median at 0.10; 25^th^ and 75^th^ percentiles at 0.08 and 0.11). The phylogenies based on Andujar’s calibration yield similar probabilities for both types of nodes (median at 0.15; 25^th^ and 75^th^ percentile at 0.14 and 0.17; and median at 0.16; 25^th^ and 75^th^ percentile at 0.14 and 0.18; respectively for pre- and post-Pleistocene nodes, see Fig. S7). These low probabilities are, nonetheless, congruent with the lack of phylogenetic signal in module co-occurrence previously found. Interestingly, and regardless of calibration scheme, the probabilities found for both type of nodes (i.e. pre- and post-Pleistocene) were higher in southern and central regions than in northern ones (Fig. 4b). This suggests that regions not covered by ice during the LGM can still reflect some old historical legacies (as shown by the higher pre-Pleistocene node probability) while accumulating some related lineages that diversify during and after the Pleistocene (as indicated by the higher post-Pleistocene node probability).

## Discussion

More than 140 years ago, Wallace (1876) foresaw that the influence of Pleistocene glaciations on the distribution of diversity had been strong enough to erode the imprint of previous events. Our results support Wallace’s thoughts, showing a remarkable coincidence between the distribution of the ice sheets at the Last Glacial Maximum and the current configuration and evolutionary structure of European *Carabus* Faunas.

The first line of evidence supporting this idea comes from the close spatial relationship between the southern limits of the ice sheet at LGM and the transition zone separating the southern and northern regions. This border also coincides with the line –identified by Calatayud et al. (2016b)– where the relationship between *Carabus* species richness and current climate changes (Fig. 2). Thus, it seems that the climate changes underwent during the Pleistocene not only shaped phylogeographic (Avise et al. 1998; Hewitt 1999; Barnes et al. 2002; Johnson et al. 2004; Theissinger et al. 2013; Horreo et al. 2016) and species richness patterns (e.g. Svenning and Skov 2007, Araújo et al. 2008, Hortal et al. 2011, Calatayud et al. 2016b), but that Ice ages have also left a strong imprint on the geographical structure of species composition at a regional scale. Accordingly, the species from the northernmost region (module 5) show the lowest level of endemism (Fig. S8), as expected for regional faunas composed of species that have recently colonized the north of Europe from southern glacial refugia (Araújo et al. 2008, Calatayud et al. 2016b, our hypothesis 5). In fact, although these species show large distribution ranges in different parts of southern Europe, their ranges only overlap near the northern Carpathian Mountains (Fig. S9). This area was a glacial refugia for a large and taxonomically diverse array of northern European species (Ursenbacher et al. 2006; Sommer and Nadachowski, 2006, Provan and Bennett 2008), including *Carabus* (Homburg et al. 2013). Additionally, the decrease in modularity values with latitude also points to a lesser geographical structure of northern assemblages, which can be interpreted as the result of a post-glacial colonization, together with less geographic complexity in some areas.

Besides the Pleistocene effects in the definition and geographical structure of regional species pools, we also found evidence of the imprint of this geological period on the processes configuring the distribution of *Carabus* faunas. The general strong relationship between regional patterns of co-occurrence and both niche similarities and spatial connectivity shows that co-occurring species tend to have similar realized environmental niches and that also tend to be geographically constrained by the same dispersal barriers. This latter result was expected given the –presumed– low dispersal capacity of *Carabus* species (see Turin et al. 2003), which is likely to be behind the spatial coincidence of module transition zones and geographical barriers. Perhaps more unexpected is the weak effect of phylogenetic distances despite the strong relationship between regional co-occurrence and niche similarities. This implies that geographical barriers rather than climatic-niche conservatism have restricted species distributions even within regions of similar climate. These results also point to that *Carabus* niche evolution is, to some extent, evolutionarily unconstrained, which is congruent with the general high adaptation capacity of insects (e.g. Overgaard and Sørensen 2008).

Whatever the origin of the relationship between species occurrence and environmental conditions, what is certainly true is that its strength changes between regions. These changes follow a latitudinal gradient in the importance of environmental niche similarities (Fig. 3). The occurrence into sub-regions is more strongly related to the similarity in the realized niche in the north than in the south. This might be a direct consequence of the effects of post-glacial colonization, where formerly glaciated areas show a clear sorting of species due to its environmental preferences. On the contrary, in southern regions, species are expected to had have more time to diversify and sort geographically by other factors besides climate (Hortal et al. 2011). Our findings corroborated this idea since we found strong effects of dispersal barriers in these areas. Moreover, although we did not find a significant phylogenetic signal in the subregional co-occurrence over these regions, our analyses revealed that they hold a small but still larger number of related species compared to northern ones, supporting that more stable regions are more prone to accumulate related species.

Despite of these related species of southern regions, we found a generalized lack of phylogenetic structuration of *Carabus* faunas. This can be the outcome of –relatively– recent speciation events due to vicariance and/or a “cul-de-sac effect” (O’Regan 2008). The former would imply the formation of dispersal barriers promoting the geographical split of many lineages and subsequent allopatric speciation (Weeks et al. 2016). Yet, the geophysical accidents that can be associated with the limits of the *Carabus* regions we found here largely predate the origin of the genus (see Beccaluva et al. 1998, Deuve et al. 2012). On the other hand, a generalized dispersion into climatic refugia, together with a subsequent stagnancy within them (i.e. a “cul-de-sac” effect) may also produce the observed mixing of unrelated linages into regions. Although it is difficult to distinguish between both processes, the latter seems more plausible, with southern regions accumulating unrelated species while acting as glacial refugia, and northern ones being recolonized by unrelated species with similar environmental niches and/or simply higher dispersal capacity (Svenning and Skov 2007).

Supporting the Pleistocene signature, our results showed a temporal coincidence between this geological period and the phylogenetic structuration of *Carabus* faunas. This result was consistent regardless of the different approaches used and across the different time calibration scenarios. This robust temporal coincidence supports that the current regional organization of *Carabus* species and lineages is rooted at the Pleistocene, which also explains the general lack of phylogenetic structure of regional *Carabus* faunas. Our results partially contrast with ancestral range estimations for clades inhabiting areas that were never glaciated, where more ancient signals were found in the spatial sorting of lineages (Condamine et al. 2015, Economo et al. 2015, Tänzler et al. 2016, Toussaint and Balke 2016). These previous findings are, nonetheless, congruent with the higher probability of holding related *Carabus* species of southern and more stable European regions. Interestingly, a visual inspection of 100 randomly chosen phylogenies (Appendix S5) reveals that several related lineages that diversified before the Pleistocene and inhabit central regions (subgenera *Orinocarabus* and *Platycarabus*; Fig. 4c and Appendix S5) mostly comprise montane and alpine species (Turing et al. 2003). This suggests that older historical signals are mostly due to presumably cold-adapted species that should be less affected by glacial conditions. Nevertheless, this speculation should be taken with caution since the central-eastern region also hosts related species (some *Morphocarabus* species, see Fig. 4) that are not particularly adapted to cold environments (Turing et al. 2003). Further studies are required to determine which characteristics allowed some *Carabus* to persist under glacial conditions. In sum, these findings suggest that the repeated advances and retreats of ice sheets and glacial conditions that characterize the European Pleistocene produced repeated cycles of retreat to southern regions and advance towards the north of *Carabus* species, a hustle-and-bustle process that ultimately led to the observed mixing of unrelated lineages, with few related species inhabiting in less affected regions.

To summarize, our results provide solid arguments in favour of the importance of Pleistocene glaciations along with geographical barriers and niche-based processes in structuring the regional faunas of European *Carabus*. On the one hand, this group’s faunas are primarily delimited by the location of the southern limit of the ice sheet at LGM, which separates two large regions that differ not only in species composition, but also in the processes underlying the spatial organization of these species. On the other hand, the phylogenetic structure of these faunas coincides with the beginning of the Pleistocene. This implies that the geographical distribution of species and lineages is profoundly shaped by past climates. Moreover, our results also suggest that ecological (Naeslund and Norberg 2006, Madrigal et al. 2016) and evolutionary mechanisms (Wüest et al. 2015, Calatayud et al. 2016a) that rely on processes occurring at regional scales can be profoundly affected by the history of Earth’s climates. Hence, the study of these historical events may be essential to unravel both large and local scale diversity patterns.

## Acknowledgements

We are very grateful to Achille Casale for providing data on *Carabus* habitat preferences and comments on an early version of the phylogeny, and the Scientific Computation Centre of Andalusia (CICA) for the computing services they provided. We acknowledge insightful discussions with Hortal lab members. JC was supported by a FPU-fellowship of the Spanish Ministry of Education (FPU12/00575). This work is partly supported by the Spanish Ministry of Economy, Industry and Competitiveness (MINECO) project SCARPO (CGL2011-29317) to JC and JH. MAR and RMV were supported through the grants CGL2017-86926-P and CGL2013-48768-P. ML and JLH were supported by MINECO FPI and Juan de La Cierva grants, respectively.

## Author contributions

JC and JH conceived research. JC designed the study with contributions of all authors. JC, RMV, ML and JLH analysed the data. All authors discussed results. JC wrote the paper with contributions of all authors.

